# Epidermal YAP Activity Drives Canonical WNT16/β-catenin Signaling to Promote Keratinocyte Proliferation *in vitro* and in the Murine Skin

**DOI:** 10.1101/278705

**Authors:** Veronica Mendoza-Reinoso, Annemiek Beverdam

## Abstract

The skin constantly self-renews throughout adult life. Wnt/β-catenin signaling plays a key role in promoting keratinocyte proliferation in the hair follicles and in the interfollicular epidermis. A recent report demonstrated that epidermal YAP activity drives β-catenin activation to promote keratinocyte proliferation in the murine skin. However, it remains unclear whether this is caused by paracrine activation of canonical Wnt signaling or through other YAP/β-catenin regulatory interactions. In the present study, we found that XAV939-inhibition of canonical WNT signaling in skin of YAP2-5SA-ΔC mice resulted in diminished β-catenin activation, reduced keratinocyte proliferation, and a mitigation of the hyperplastic abnormalities in the interfollicular epidermis, signifying a canonical WNT ligand-dependent mechanism. Our subsequent analyses determined that WNT16 is produced in response to YAP activity in keratinocytes both *in vitro* and *in vivo*, and that WNT16 drives HaCaT keratinocyte proliferation via canonical WNT16/β-catenin signaling. We conclude that under normal physiological conditions WNT16 is the paracrine WNT ligand secreted in response to epidermal YAP activity that promotes cell proliferation in the interfollicular epidermis. This study delineates a fundamental YAP-driven mechanism that controls normal skin regeneration, and that may be perturbed in human regenerative disease displaying increased YAP and WNT signaling activity.

## INTRODUCTION

The skin is a unique organ characterized by its continuous regeneration. In order to maintain normal skin homeostasis, epidermal stem/progenitor cell proliferation must be tightly regulated, and any variation of this well-balanced process may result in the development of skin disease. However, the molecular mechanisms involved in epidermal stem/progenitor cell proliferation remain poorly understood.

The Hippo kinase pathway controls organ development, tissue regeneration and stem/progenitor cell self-renewal (Camargo et al., 2007; Dong et al., 2007; Lian et al., 2010). This kinase cascade is classically known to phosphorylate the downstream effectors Yes-Associated Protein (YAP) and the transcriptional co-activator with PDZ-binding motif (TAZ), resulting in their inactivation through cytosolic retention. Unphosphorylated YAP/TAZ translocate to the nucleus and bind the Transcriptional Enhancer Associate Domain (TEAD) transcription factors to activate gene expression and cell proliferation. Recently, mechanical factors and the tissue microenvironment were shown to play a major overarching role over the Hippo kinase pathway in controlling YAP/TAZ activity, which since then are known to act as mechanosensors in control of tissue homeostasis (Aragona et al., 2013; Dupont et al., 2011; Piccolo, Dupont, & Cordenonsi, 2014; Yu, Zhao, & Guan, 2015).

The canonical WNT pathway also has a crucial role during development, cell fate determination and stem/progenitor self-renewal in many tissues (Clevers, 2006). WNT signaling is activated by binding of WNT ligands to the Frizzled transmembrane receptors. This results in the dissociation of β-catenin from the cytosolic destruction complex, and its nuclear translocation and binding to TCF/LEF family transcription factors to activate the expression of target genes and epidermal stem/progenitor cell proliferation (Clevers, Loh, & Nusse, 2014; Huelsken, Vogel, Erdmann, Cotsarelis, & Birchmeier, 2001; Lo Celso, Prowse, & Watt, 2004). Non-canonical WNT pathways such as the planar cell polarity and WNT/calcium pathways act independently of the cytosolic destruction complex and β-catenin transcriptional activity, and mainly control cellular polarity and motility (Katoh, 2005; Kohn & Moon, 2005).

Canonical WNT/β-Catenin signaling has well-established roles in the activation of keratinocyte proliferation during hair follicle cycling (Huelsken et al., 2001). Furthermore, recent reports have resolved a longstanding contentious issue in the field, and unequivocally established that paracrine Wnt signaling and β-Catenin transcriptional activity are required to promote basal keratinocyte proliferation in the interfollicular epidermis (Choi et al., 2013; Lim et al., 2013), and not keratinocyte differentiation, as initially thought (Huelsken et al., 2001). However, the WNT ligands that are responsible for driving β-Catenin transcriptional activity in the interfollicular epidermis remain unknown.

The Hippo/YAP pathway interacts with many signaling pathways to control the homeostasis of tissues, including in the skin (Akladios, Mendoza-Reinoso, et al., 2017; Akladios, Mendoza Reinoso, et al., 2017; Piccolo et al., 2014). Cross-talk between the Hippo/YAP and WNT/β-Catenin signaling pathways was reported to be mediated by the cytosolic β-catenin destruction complex, through regulatory interactions taking place in the cell nuclei (Heallen et al., 2011; Rosenbluh et al., 2012; Wang et al., 2014), or through other mechanisms (Azzolin et al., 2014; Cai, Maitra, Anders, Taketo, & Pan, 2015; Imajo, Miyatake, Iimura, Miyamoto, & Nishida, 2012; Oudhoff et al., 2016; Park & Jeong, 2015). Recently, interactions between these two pathways were also shown to occur during epidermal homeostasis, and we and others showed that increased YAP activity in the basal keratinocytes of murine skin results in β-catenin activation (Akladios, Mendoza-Reinoso, et al., 2017), and in severe epidermal hyperplasia in the interfollicular epidermis and in the hair follicles (Beverdam et al., 2013; Schlegelmilch et al., 2011; Zhang, Pasolli, & Fuchs, 2011). However, the precise underlying molecular mechanism of how epidermal YAP activity promotes β-catenin activity to drive keratinocyte proliferation remains elusive.

In the present study, we found that inhibition of canonical Wnt/β-catenin signaling in the skin of YAP2-5SA-ΔC mice results in an amelioration of the epidermal hyperplasia in the interfollicular epidermis. In addition, we found that epidermal YAP promotes WNT16 expression in HaCaT keratinocytes and in the mouse interfollicular epidermis *in vivo*. Furthermore, we established that WNT16 promotes keratinocyte proliferation both independently of β-catenin, and through canonical WNT16/β-catenin signaling depending on the levels of WNT16 expression. We conclude that WNT16 is the WNT ligand that under normal physiological conditions promotes canonical WNT/β-catenin signaling to drive proliferation of keratinocytes in response to epidermal YAP activity *in vitro* and in the murine interfollicular epidermis *in vivo*.

## MATERIAL AND METHODS

### Animals

All animal experimental procedures performed were conducted under protocols approved by the UNSW Australia’s Animal Care and Ethics Committee Unit, and in compliance with the National Health and Medical Research Council ‘Australian code of practice (8^th^ edition, 2013). Thirty-eight days old YAP2-5SA-ΔC mice were topically treated with 100μl of 5μM XAV-939 (Chem-Supply, X0077-25MG) diluted in DMSO and vehicle solution (DMSO) daily for 13 days.

### Tissue processing and Histological and Immunofluorescence staining

Full thickness mice skin tissues were fixed in 4% paraformaldehyde, paraffin embedded, sectioned, and histology-stained following routine procedures. Antigen retrieval was performed using 10mM sodium citrate buffer (pH 6.0) and a Milestone RHS-1 Microwave at 110^°^C for 5 minutes. Tissue sections were immunostained using previously standardized methods, and confocal images were captured using an Olympus FV1200 laser scanning confocal microscope. Epidermal thickness and immuno-signal intensity were quantified using 3 mice and 3 skin regions per mouse. Immuno-signal was quantified in a semi-automated fashion using ImageJ software. Antibody information is available in Table S1.

### Western blots and qRT-PCR analysis

Protein and RNA were isolated from full thickness mouse skin biopsies and HaCaT cells using TRIzol^®^ reagent (Thermo Fisher Scientific, 15596026) following standard protocols. Protein lysates were analysed by western blot, intensity of bands was quantified with ImageJ software and normalized to GAPDH, GSK3β or β-catenin. Primary and secondary antibody information is available in Table S1. Quantitative RT–PCR assays were carried out using Fast SYBR^®^ Green Master Mix (Thermo Fisher Scientific, 4385612) and Mx3000P qPCR System (Agilent Technologies), and were analysed by the comparative cycle time method, normalizing to *18S* ribosomal or *GAPDH* RNA levels. Human and mouse primer information is available in Table S2.

### Cell lines and transfections

HaCaT immortalized keratinocytes of passages between 15 and 21 were maintained in DMEM/F-12 (Sigma, D8062), supplemented with 10% FBS (Gibco, 10437-028) and 1X Penicillin-Streptomycin (Gibco, 15140-122) in a 5% CO2 incubator at 37°C. Transient transfections were performed using Lipofectamine3000 (Thermo Fisher Scientific, L3000015) according to manufacturer’s instructions. To overexpress dominant active YAP, pDsRed Monomer C1-YAP2-S127A mutant (Addgene, 19058) and pDsRed Monomer vector control DNA were used. WNT16 overexpression was carried out using pcDNA3.2-WNT16-V5 (Addgene, 35942) and pcDNA3.2 vector control. *YAP* knockdown was performed using MISSION^®^ Universal and *YAP* siRNA (Sigma). *WNT16* knockdown was performed using MISSION^®^ *GFP* and *WNT16* esiRNA (Sigma). MTT assays were performed using Thiazolyl Blue Tetrazolium Bromide (Sigma). Immunostaining assays were performed following standard protocols. Antibody information is available in Table S1. Imaging was performed using an Olympus FluoView^™^ FV1200 Confocal Microscope.

### Statistical analysis

Statistical significance was determined by Student’s unpaired t-tests. Error bars represent mean ± SEM. Asterisks indicate statistical significance, where *P* < 0.05 was used as significance cut-off.

## RESULTS

### XAV939 inhibition of canonical WNT signaling ameliorates epidermal hyperplasia of YAP2-5SA-ΔC mice

We recently established that epidermal YAP activity promotes β-catenin activity to drive keratinocyte proliferation in the mouse skin *in vivo* (Akladios, Mendoza-Reinoso, et al., 2017). However, it remains unclear whether these interactions are mediated by paracrine activation of canonical Wnt signaling, or by other YAP/β-catenin regulatory interactions. Firstly, we assessed whether these were caused by activation of canonical Wnt/β-catenin signaling activity. To do so, we used YAP2-5SA-ΔC mice, which display increased epidermal YAP activity, increased β-catenin activity, and severe hyperplasia of the basal layer due to increased keratinocyte proliferation. Furthermore, the skin of these mice also display abnormal hair follicles and hyperkeratosis (Akladios, Mendoza-Reinoso, et al., 2017; Beverdam et al., 2013). We topically treated YAP2-5SA-ΔC mice with XAV-939 or vehicle daily for thirteen days. XAV-939 is a small molecule that selectively inhibits canonical (and not non-canonical) Wnt signaling through stabilizing Axin2 and promoting the destruction of cytosolic β-catenin (Distler et al., 2013).

Histological analysis showed that the thickness of the hyperplastic interfollicular epidermis of XAV-939-treated YAP2-5SA-ΔC mouse skin was approximately a third of that of vehicle-treated YAP2-5SA-ΔC mouse skin (Figure 1c-d.1; *P*<0.04, *N*=3). Effective inhibition of Wnt/β-catenin signaling activity by XAV-939 was confirmed by reduced nuclear active β-catenin (Figure 1f, f.1; *P*<0.002, *N*=3), and reduced nuclear localization of β-catenin direct target Cyclin D1 (Figure 1h, h.1; *P*<0.0001, *N*=3) in the interfollicular epidermis. Furthermore, nuclear expression of the basal epidermal stem/progenitor cell marker P63 (*P*<0.0001, *N*=3) (Figure 1i *vs.* j & j.1), and of proliferation marker Ki67 (*P*<0.008, *N*=3) (Figure 1k *vs.* l & l.1) were also significantly reduced in the XAV-939- *vs.* vehicle-treated YAP2-5SA-ΔC interfollicular epidermis. Altogether, these data are in line with our previous findings (Akladios, Mendoza-Reinoso, et al., 2017), and furthermore establish that epidermal YAP activity promotes canonical Wnt signaling to drive β-catenin nuclear activity and basal keratinocyte proliferation in the interfollicular epidermis of the mouse skin *in vivo*.

**Figure 1.**
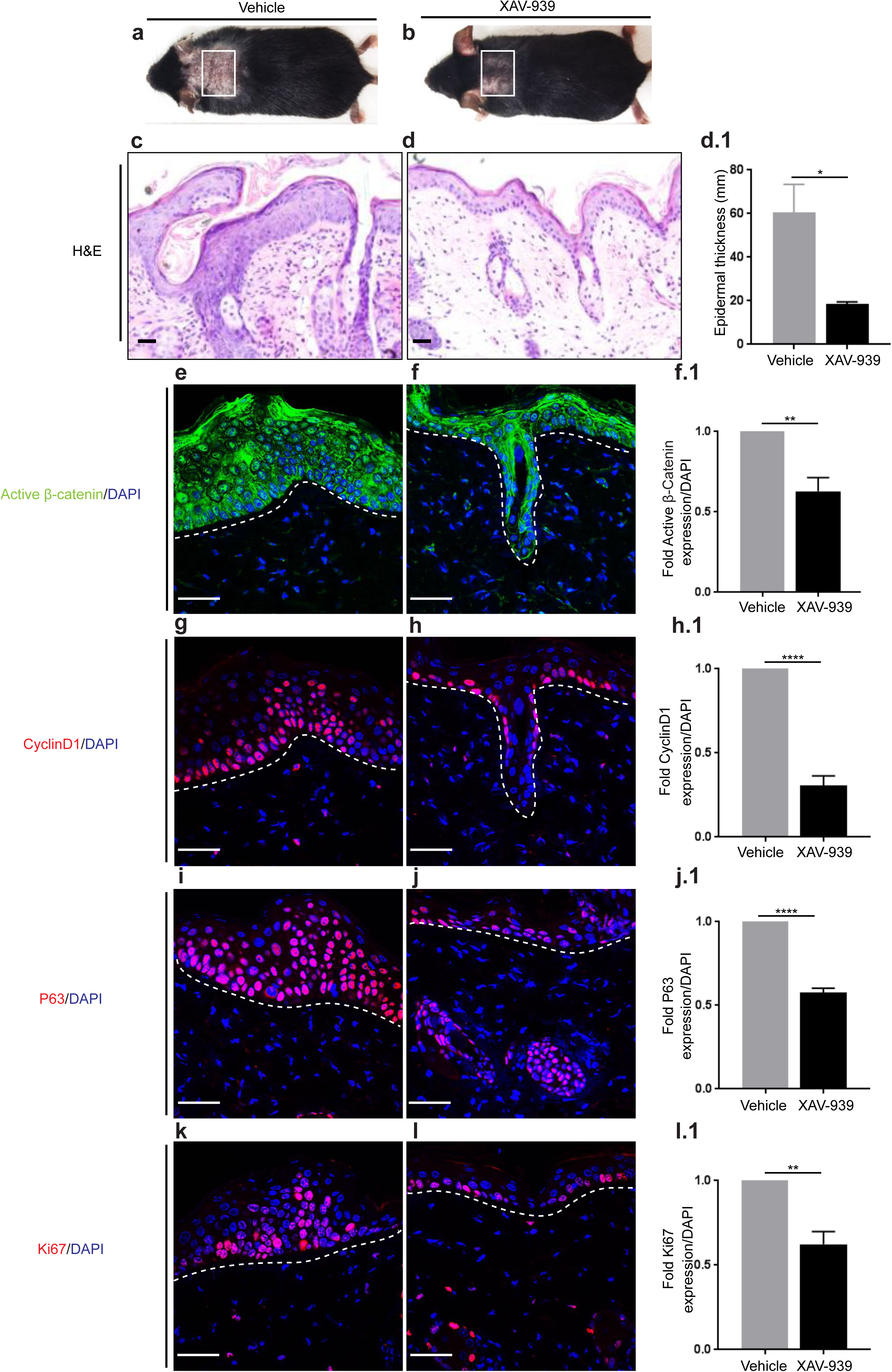
XAV939 inhibition of canonical WNT signaling ameliorates epidermal hyperplasia of YAP2-5SA-ΔC mice. Representative images of YAP2-5SA-ΔC mice treated with vehicle (a, c, e, g, i, k), or XAV-939 (b, d, f, h, j, l). H&E staining of boxed skin areas (c,d). Quantification of epidermal thickness (d.1). Immunofluorescence analysis showing active β-catenin (e,f), Cyclin D1 (g,h), p63 (i, j) and Ki67 (k, l) staining. Quantification of active β-catenin (f.1), Cyclin D1 (h.1), p63 (j.1) and Ki67 (l.1) positive nuclei in the epidermis. H&E, Hematoxylin and eosin; DAPI, 4, 6-diamidino-2-phenylindole. Scale bars = 20 *μ*m.

### YAP activity promotes *WNT16* expression in keratinocytes grown *in vitro* and in the murine epidermis

Next, we wanted to identify which WNT ligands are responsible for driving WNT/β-catenin signaling in response to epidermal YAP activity to promote keratinocyte proliferation. Transcriptomics approaches found that *Wnt3, Wnt3a, Wnt4, Wnt6, Wnt7A, Wnt7B, Wnt10A, Wnt10B* and *Wnt16* are expressed in the developing or regenerating mouse epidermis (Rezza et al., 2016; Sennett et al., 2015). These and their human orthologues were selected for analyses. Quantitative real time RT-PCR assays revealed significantly increased expression of *Wnt16* expression (Figure 2a. *P*<0.0001, *N*=3), and significantly decreased expression of *Wnt3, Wnt3a, Wnt4, Wnt6* and *Wnt10b* (Figure 2a, *P*<0.0025, *N*=3) in YAP2-5SA-ΔC *vs*. wildtype skin. Then we assessed the expression of these *WNT* ligand genes upon YAPS127A overexpression in immortalized human HaCaT keratinocytes. We chose to overexpress YAPS127A rather than YAP2-5SA-ΔC because previous reports have shown that the latter acts as a dominant negative YAP mutant protein *in vitro* due to loss of its transactivation domain encoded by its C-terminus (Hoshino et al., 2006; Zhao et al., 2007). We found significantly increased gene expression of *YAP* (*P*<0.0005; *N*=3), and of YAP/TEAD direct target genes *CTGF* (*P*<0.0002; *N*=3) and *ANKRD1* (*P*<0.04; *N*=3) in YAPS127A *vs*. control-transfected HaCaT keratinocytes, confirming YAP activation in YAPS127A-transfected keratinocytes (Figure 2b). Furthermore, expression of *WNT16* (*P*<0.007; *N*=3) was also increased, whereas *WNT6* (*P*<0.007; *N*=3) and *WNT10B (P*<0.03; *N*=3) expression levels were decreased in YAPS127A *vs*. control-transfected HaCaT keratinocytes (Figure 2b). *WNT3, WNT3A, WNT4, WNT7A, WNT7B*, and *WNT10A* expression levels were unchanged (*N*=3, Figure 2b). si*YAP*-transfected HaCaT keratinocytes on the other hand, displayed decreased *YAP* expression levels (*P*=0.0002; *N*=3), confirming effective knock down, and reduced expression of *WNT4* (*P*=0.0007; *N*=3), *WNT7B* (*P*<0.008; *N*=3), *WNT10B* (*P*=0.03; *N*=3), and *WNT16* (*P*<0.007; *N*=3) compared to scramble control-transfected cells (Figure 3c). *WNT10A* expression levels were slightly increased (*P*<0.005; *N*=3) (Figure 3c), whereas expression of *WNT3, WNT3A, WNT6* and *WNT7A* were unchanged (*N*=3; Figure 3c). These data show that YAP activity positively controls the expression levels of *WNT16* in proliferating HaCaT keratinocytes grown *in vitro*.

**Figure 2.**
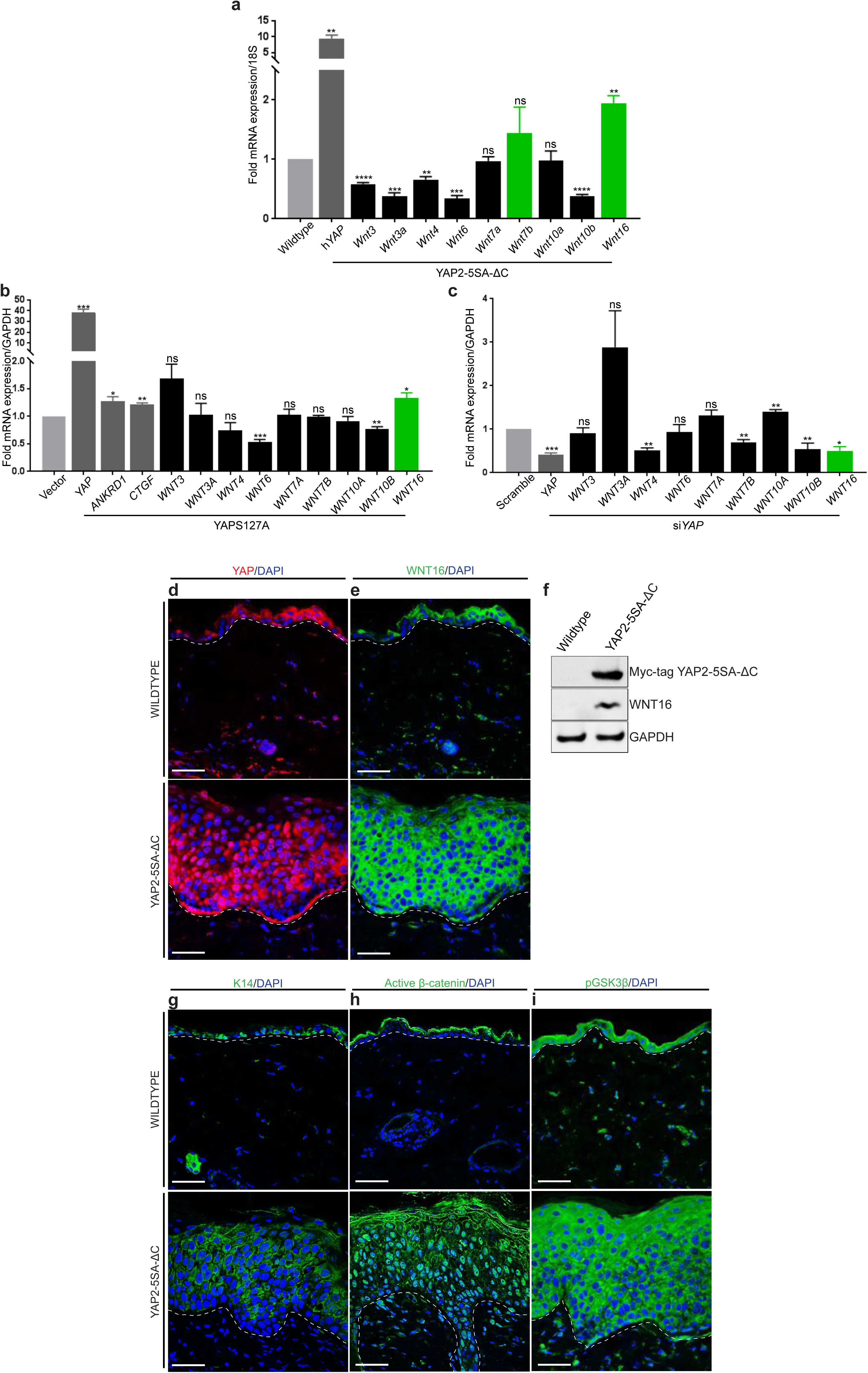
YAP activity promotes *WNT16* expression in keratinocytes grown *in vitro* and in the murine epidermis. Quantitative RT-PCR analyses showing *WNT3, WNT3A, WNT4, WNT6, WNT7A, WNT7B, WNT10A, WNT10B* and *WNT16* expression levels relative to 18S or *GAPDH* in lysates of YAP2-5SA-ΔC skin (a) and HaCaT keratinocytes transfected with YAPS127A (b) or si*YAP* (c). Immunofluorescence staining of dorsal skin sections of adult wildtype and YAP2-5SA-ΔC littermate mice detecting K14 (e), YAP (f), active β-catenin (g), WNT16 (h), phospho-GSK3β (i). DAPI, 4, 6-diamidino-2-phenylindole. Scale bars = 20 *μ*m.

**Figure 3.**
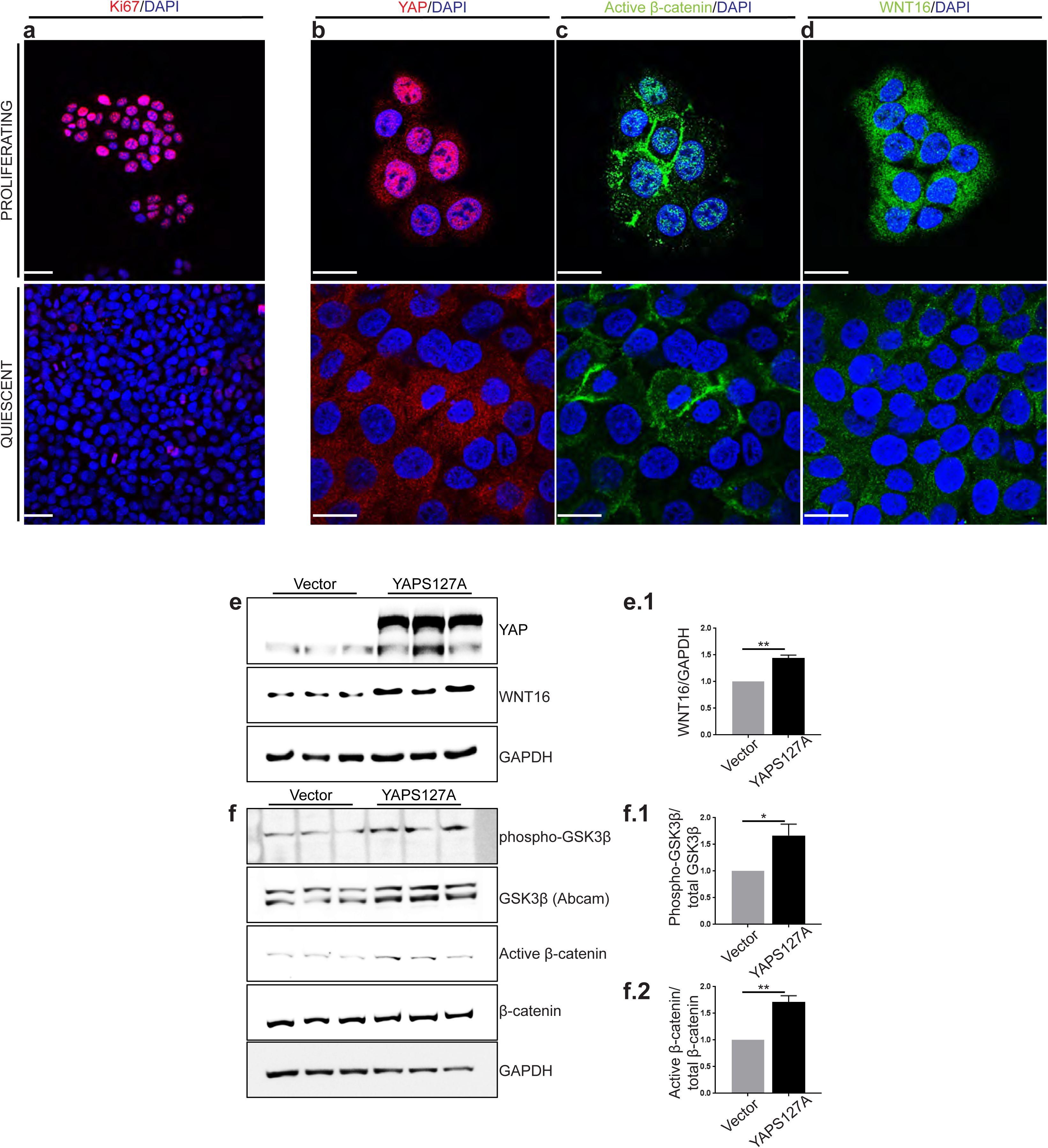
YAP activity drives canonical WNT16/β-catenin signalling in proliferating keratinocytes. (a-d) Immunofluorescence staining of proliferating (top) and quiescent (bottom) HaCaT keratinocytes showing Ki67 (a), YAP (b), active β-catenin (c) and WNT16 expression (d). Western blots of (e) YAP, WNT16, GAPDH, and (f) phospho-GSK3β, total GSK3β, active β-catenin, total β-catenin, and GAPDH, in protein lysates of vector and YAPS127A transfected HaCaT keratinocytes (*N*=3). Quantification of WNT16 expression (e.1), GSK3β phosphorylation (f.1), and active β-catenin expression (f.2).

Next, we investigated gene and protein expression in skin biopsies of YAP2-5SA-ΔC mice, and first confirmed significantly higher *hYAP* (*P*<0.003; *N*=3) and *Wnt16* (*P*<0.02; *N*=3) RNA expression levels compared to in skin biopsies of wildtype littermate mice (data not shown). Moreover, immunofluorescence assays displayed increased nuclear YAP and increased activated β-catenin in the dramatically thickened basal Keratin14-expressing layer of the interfollicular epidermis of YAP2-5SA-ΔC mice compared to wildtype (Figure 2d, g, h), which is in line with previously reported (Akladios, Mendoza-Reinoso, et al., 2017; Beverdam et al., 2013). Furthermore, the YAP2-5SA-ΔC basal epidermis displayed increased WNT16 protein levels (Figure 2e, f), and increased levels of phospho-GSK3β relative to wildtype littermate skin (Figure 3i). These results show that YAP activity in basal keratinocytes promotes WNT16 expression and canonical WNT/β-catenin signaling in the regenerating mouse interfollicular epidermis *in vivo*.

### YAP activity in proliferating keratinocytes promotes canonical WNT16/β-catenin signaling

A previous study reported that WNT16B overexpression in normal human keratinocytes (NHEKs) resulted in activation of non-canonical WNT signaling and increased cell proliferation, but not in β-catenin activation (Teh et al., 2007). However, WNT16 is reported to act both as a canonical and as a non-canonical WNT ligand (Christodoulides, Lagathu, Sethi, & Vidal-Puig, 2009; Mazieres et al., 2005; Moverare-Skrtic et al., 2014; Nalesso et al., 2017).

To understand if YAP activity promotes canonical WNT16/β-catenin signaling to drive keratinocyte proliferation, immunofluorescence assays were performed in HaCaT keratinocytes grown *in vitro* at low and high density. Predominantly nuclear YAP and activated β-catenin were detected in low-dense proliferating keratinocytes as previously reported (Akladios, Mendoza-Reinoso, et al., 2017). Furthermore, WNT16 protein expression levels were relatively high compared to in high dense and quiescent keratinocytes in which YAP and β-catenin were sequestered in the cytosol and the cell junctions (Figure 4a-d). These data are consistent with the hypothesis that WNT16 drives β-catenin activation and cell proliferation in response to YAP activity in keratinocytes.

**Figure 4.**
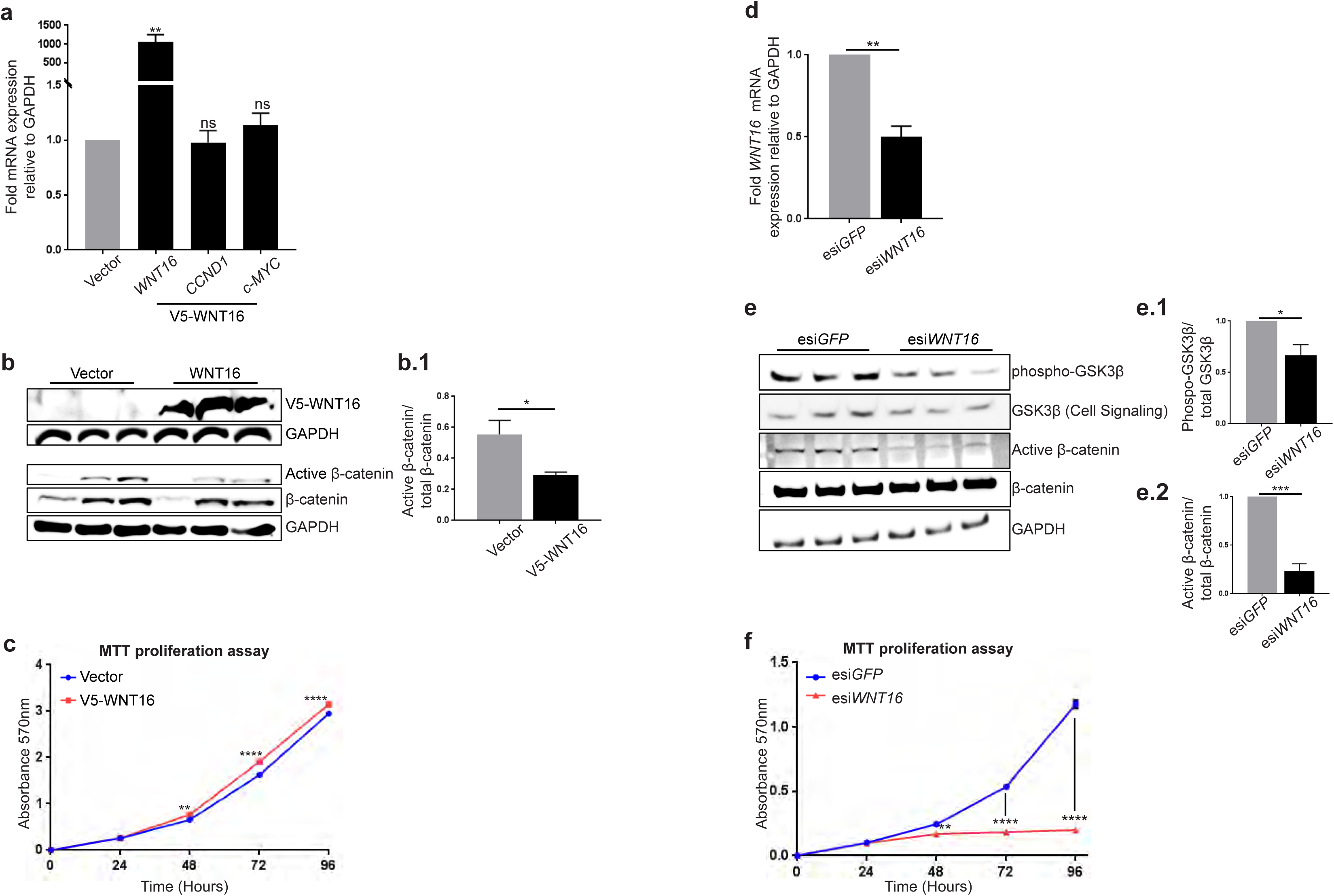
WNT16 promotes cell proliferation independently of β-catenin activation or through canonical WNT16/β-catenin signaling. (a) Fold-change of *WNT16, CCND1* and *cMYC* expression normalized to *GAPDH* in HaCaT keratinocytes transfected with V5-WNT16 or vector control. (b) Western blots detecting V5-WNT16, active β-catenin, total β-catenin, and GAPDH in protein lysates of vector control and V5-WNT16 (right) transfected HaCaT keratinocytes (*N*=3). (b.1) Quantification of fold-change protein expression of active β-catenin normalized to total β-catenin lysates of vector control and V5-WNT16 transfected HaCaT keratinocytes. (c) MTT proliferation assay using HaCaT keratinocytes transfected with vector control and V5-WNT16 (*N*=3). Fold-change of *WNT16* mRNA expression normalized to *GAPDH* in HaCaT keratinocytes transfected with esi*GFP* and esi*WNT16* RNA (d). Western blots showing phospho-GSK3β expression normalized to total GSK3β (e and e.1) and active β-catenin normalized to total β-catenin (e and e.2) in protein lysates of esi*GFP*- and esi*WNT16*-transfected (right) HaCaT keratinocytes. (f) MTT proliferation assay using HaCaT keratinocytes transfected with esi*GFP* and esi*WNT16* RNA (*N*=3).

To further test this hypothesis, we assessed the effect of increased YAP activation on WNT16 expression levels, GSK3β phosphorylation, and β-catenin activation in HaCaT keratinocytes. WNT16 protein expression was indeed increased in YAPS127A-transfected cells (*P*<0.015; *N*=3; Figure 4e, e.1), in line with our RNA expression data (Figure 2a). Furthermore, we observed increased phospho-GSK3β (*P*<0.04; *N*=3; Figure 4f, f.1), and increased activated β-catenin (*P*<0.004; *N*=3; Figure 4f, f.2) compared to control-transfected cells. These data show that YAP activity increases WNT16 expression and canonical WNT/β-catenin signaling in proliferating HaCaT keratinocytes *in vitro*.

### WNT16 promotes keratinocyte proliferation through canonical WNT16/β-catenin signaling

Next, we tested the effect of WNT16 overexpression on β-catenin activation and cell proliferation in HaCaT keratinocytes. *WNT16* RNA and WNT16 protein expression levels were increased in V5-WNT16- *vs.* control transfected HaCaT keratinocytes (Figure 4a and b), levels of activated *vs.* pan β-catenin were significantly reduced (Figure 4b and b.1), and expression levels of β-catenin transcriptional targets *CCDN1* and *cMYC* were unchanged (Figure 4a). Furthermore, V5-WNT16-transfected HaCaT keratinocytes proliferated at a mildly but significantly increased rate (Figure 4c). These data confirm that WNT16 overexpression promotes keratinocyte proliferation, but not β-catenin activity, in line with the earlier study (Teh et al., 2007).

A recent publication showed that WNT16, in the presence of excessive levels of WNT ligands, antagonised canonical WNT signaling (Nalesso et al., 2017). So conceivably, the high levels of WNT16 expression in overexpression studies with NHEK (Teh et al., 2007) and HaCaT keratinocytes (this study) may have caused a shutdown of canonical WNT signaling in these cells. To test the effect of reduced *WNT16* expression on canonical WNT signaling and keratinocyte proliferation, we assessed GSK3β phosphorylation, β-catenin activity, and cell proliferation in esi*Wnt16*- *vs.* control transfected HaCaT keratinocytes. Quantitative real time RT-PCR assays confirmed reduced *WNT16* RNA expression levels in esi*WNT16*- *vs*. control-transfected keratinocytes (*P*<0.002; *N*=3; Figure 4d). Furthermore, protein expression levels of phospho-GSK3β (*P*<0.05; *N*=3; Figure 4e and e.1), and of activated β-catenin (*P*<0.0005; *N*=3; Figure 4e and e.2) were reduced in esi*WNT16*-transfected HaCaT keratinocytes, and MTT assays established significantly reduced cell proliferation rates in esi*WNT16*-*vs*. control transfected HaCaT keratinocytes (Figure 4f). These data show that at physiological levels WNT16 promotes proliferation of HaCaT keratinocytes through activation of canonical WNT16/β-catenin signaling.

Altogether, these data establish that YAP activity in keratinocytes promotes WNT16 expression and keratinocyte proliferation via canonical WNT16/β-catenin signaling.

## DISCUSSION

Recent investigations have demonstrated that YAP drives keratinocyte proliferation in the murine epidermis *in vivo* and in HaCaT keratinocytes *in vitro* (Akladios, Mendoza-Reinoso, et al., 2017; Beverdam et al., 2013; Schlegelmilch et al., 2011; Zhang et al., 2011) through promoting β-catenin activity (Akladios, Mendoza-Reinoso, et al., 2017). However, it is still unclear whether YAP drives β-catenin activation and keratinocyte proliferation through paracrine activation of canonical Wnt signaling, or through other regulatory mechanisms. Here, using two independent experimental models, we demonstrate that epidermal YAP activity drives WNT16 expression to promote keratinocyte proliferation through canonical WNT16/β-catenin signaling (Figure 5a). These findings are consistent with reports showing that WNT16 promotes keratinocyte proliferation *in vitro* (Teh et al., 2007), and in the murine hair follicle bulge *in vivo* (Kandyba et al., 2013).

**Figure 5.**
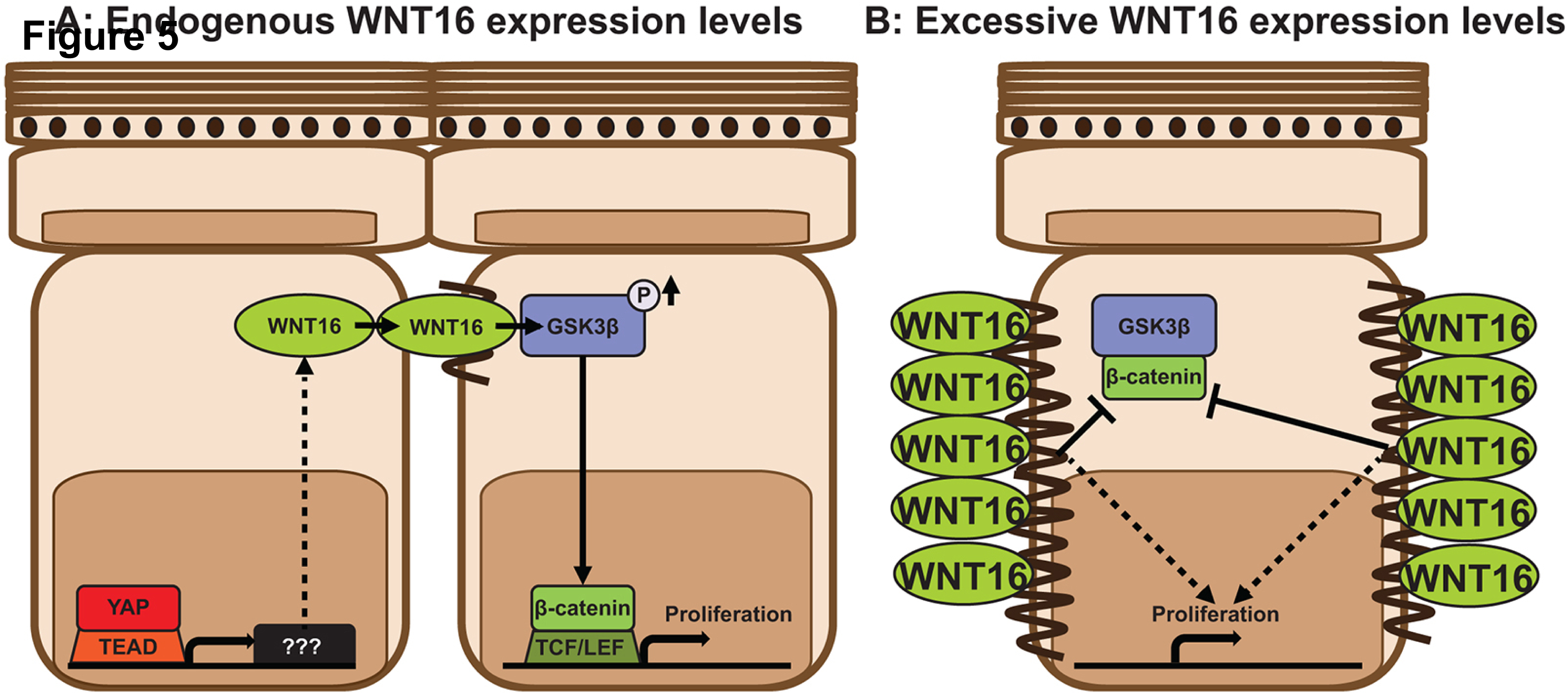
YAP promotes WNT16/β-catenin signalling to drive keratinocyte proliferation. (a) Working model of how YAP activity in keratinocytes promotes WNT16 production and canonical WNT/β-catenin signalling to drive cell proliferation in the murine epidermis. (b) Excessive WNT16 signals drive keratinocyte proliferation independently of β-catenin activity.

Recent reports resolved a contentious issue on the role of β-catenin in the control of keratinocyte proliferation in the interfollicular epidermis, and unequivocally demonstrated that auto/paracrine WNT/β-catenin signaling promotes keratinocyte proliferation, and not differentiation as suggested by earlier studies (Choi et al., 2013; Huelsken et al., 2001; Lim et al., 2013). However, the responsible WNT ligands remained elusive. Our studies are in support of these two recent reports, and furthermore reveal WNT16 as a WNT ligand driving canonical WNT/β-catenin signaling and keratinocyte proliferation *in vitro* and *in vivo*.

WNT16 is generally recognized to act both as a canonical and as a non-canonical WNT ligand in osteoblasts (Moverare-Skrtic et al., 2014), chondroblasts (Nalesso et al., 2017), leukemia (Mazieres et al., 2005), and in adipogenesis (Christodoulides et al., 2009). A previous study reported that WNT16B overexpression in normal human keratinocytes (NHEKs) results in increased cell proliferation, but not in increased canonical WNT signaling or β-catenin activation (Teh et al., 2007). Instead, it was reported to activate a non-canonical WNT signaling pathway through the N-Jun-terminal kinase cascade *in vitro* (Teh et al., 2007). Our studies confirm that WNT16 positively regulates keratinocyte proliferation. This was very obvious from our studies in which we repressed WNT16 expression in HaCaT keratinocytes (Figure 4f) relative to those where we overexpressed WNT16 in HaCaT keratinocytes (Figure 4c). The relatively small effect of WNT16 overexpression on promoting keratinocyte proliferation may be explained by the fact that keratinocytes were already in a highly proliferative state and were unable to be induced even further. We also found that WNT16 overexpression did not result in β-catenin activation in HaCaT keratinocytes. Rather, it resulted in reduced β-catenin activation in these cells (Figure 5b). Nevertheless, we also showed that inhibition of endogenous WNT16 expression in HaCaT keratinocytes by siRNAs resulted in decreased canonical Wnt signaling and β-catenin activity, and reduced cell proliferation, demonstrating that WNT16 can also promote canonical Wnt signaling to drive keratinocyte proliferation. These seemingly contradictory observations may be explained by a recent publication, which proposed that WNT16 will counteract canonical WNT signaling in the presence of excessive levels of WNT ligands (Nalesso et al., 2017). Since in our RNA interference studies, we repressed endogenous levels of WNT16 expression, we propose that under normal physiological conditions WNT16 will act as a canonical WNT ligand to promote keratinocyte proliferation (Figure 5a).

In contrast, WNT16 protein expression was mildly elevated due to increased YAP activity in the YAPS127A-transfected HaCaT keratinocytes (Figure 3e-f) and in the YAP2-5SA-ΔC epidermis (Figure 2d-h). Since we also detected concurrent activation of canonical WNT signaling, we conclude that the WNT16 expression levels induced by YAP in these systems drive cell proliferation, but are apparently not high enough to lead to repression of canonical WNT signaling through this recently reported counteracting mechanism (Nalesso et al., 2017). Interestingly, the mildly increased epidermal WNT16 expression levels in skin of YAP2-5SA-ΔC mice still appear to be somewhat compensated for by means of reduced expression levels of other WNTs expressed in the skin biopsies (Figure 2a) through an alternative unknown mechanism.

How YAP activates *WNT16* expression remains unclear. We found that *WNT16* expression is tightly regulated by YAP, indicating that it may be a direct YAP/TEAD transcriptional target in basal keratinocytes. However, a previous study in breast cancer cells failed to identify *WNT16* as a direct YAP/TEAD target gene in a different epithelial cell type (Zanconato et al., 2015). Therefore, *WNT16* expression may also be activated indirectly, in response to other effects downstream of YAP/TEAD transcriptional activity in epithelial cells (Figure 5a).

There is a heightened interest in the stem cell field with an ever-increasing body of high impact publications demonstrating that the Hippo/YAP and WNT/β-catenin pathways interact to control cell proliferation in tissue homeostasis and in cancer development. A recent study proposed a model where YAP/TAZ are integral components of β-catenin destruction complex, which explained a number of contradictory published observations (Azzolin et al., 2014; Azzolin et al., 2012; Oudhoff et al., 2016). In this model, the nuclear translocation of YAP and β-catenin depends on the presence of WNT ligands, and the activation of canonical WNT signaling. In our study, we established that YAP activity promotes WNT16 expression in epidermal regeneration, revealing that YAP may be able to control its own activity by promoting the dissociation of the β-catenin destruction complex. Our work may therefore reveal yet another level of regulatory interactions between YAP and β-catenin in stem cell control during tissue homeostasis, but does not exclude that these proteins may also undergo additional regulatory interactions to control epidermal stem/progenitor cell proliferation, for instance similar to those that were previously reported to occur in the nucleus (Heallen et al., 2011; Rosenbluh et al., 2012; Wang et al., 2014), in the β-catenin destruction complex (Azzolin et al., 2014; Cai et al., 2015), or otherwise (Imajo et al., 2012; Oudhoff et al., 2016; Park & Jeong, 2015).

Uncontrolled proliferation of tissue stem/progenitor cells can result in regenerative disease such as cancer. Interestingly, both YAP and WNT signaling have been implicated in carcinogenesis in various tissues (reviewed in Piccolo et al., 2014), including in the skin (reviewed in Andl, Zhou, Yang, Kadekaro, & Zhang, 2017). Furthermore, YAP/TAZ have been reported to regulate mesenchymal and skeletal stem cells during osteogenesis (Tang, Feinberg, Keller, Li, & Weiss, 2016; Tang & Weiss, 2017), and WNT16 plays a well-recognized role in joint formation and bone homeostasis (Guo et al., 2004; Moverare-Skrtic et al., 2014; Wergedal, Kesavan, Brommage, Das, & Mohan, 2015). Therefore, our findings have important implications for our understanding of the molecular aetiology of regenerative skin disorders, cancer, and cartilage and joint diseases that display increased nuclear YAP and/or WNT16 activity.

## ACKNOWLEDGMENTS

We thank the members of the Developmental and Regenerative Dermatology Laboratory and Dr. Nicole Bryce for helpful feedback on the project, and Dr. Megan Wilson (University of Otago) for critically reading our manuscript drafts. We thank the ABR (Garvan) and BRC (UNSW Sydney) for support with animal experimentation. We thank the BMIF (UNSW Sydney) for imaging support. This work was supported by the National Health and Medical Research Council of Australia. Miss Mendoza-Reinoso is a recipient of Tuition Fee Scholarship by UNSW Australia and Doctorate Overseas Scholarship by FINCyT Peru.

## FIGURE LEGENDS

**Table S1.**
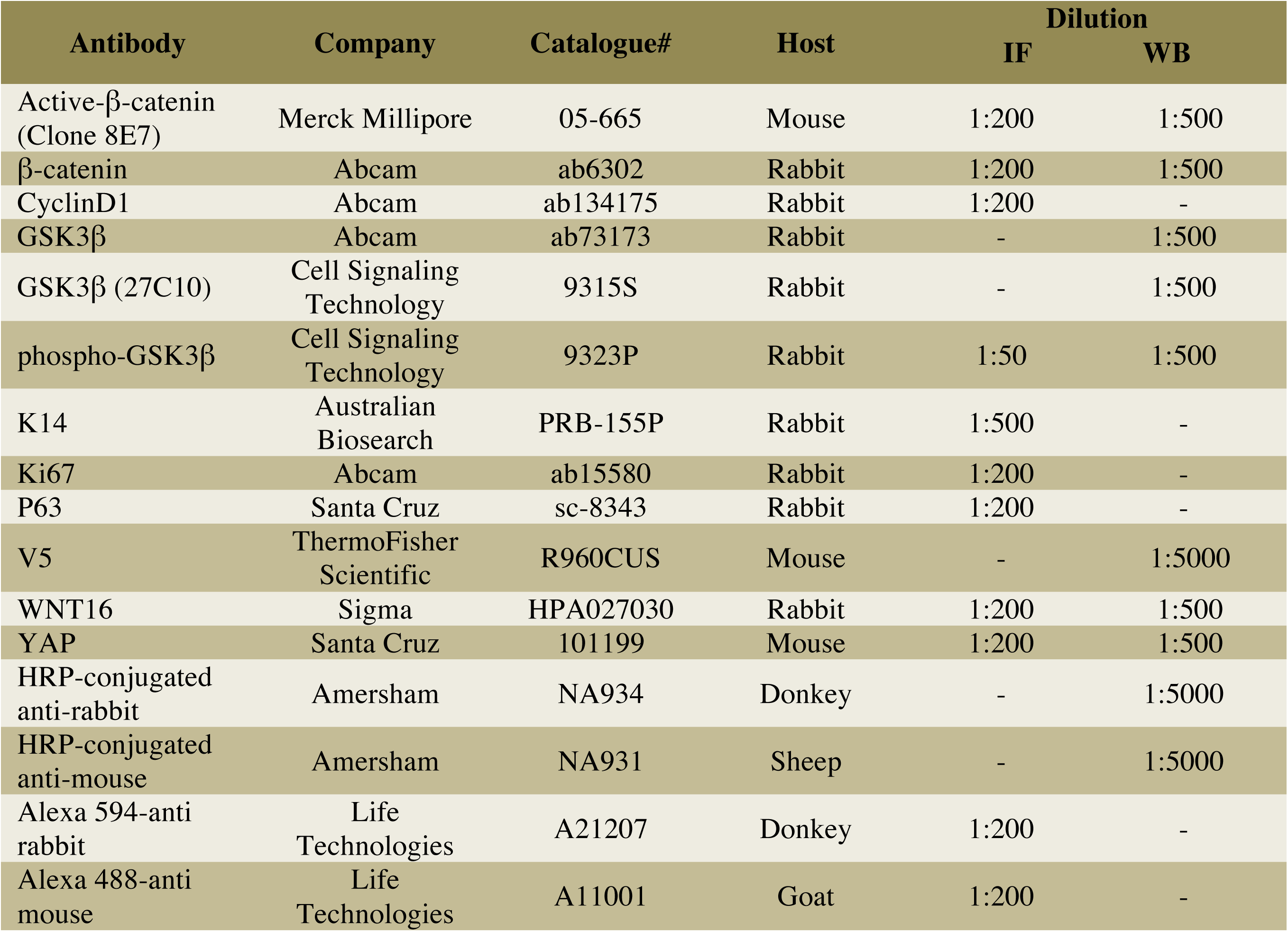
List of primary and secondary antibodies.

**Table S2.**
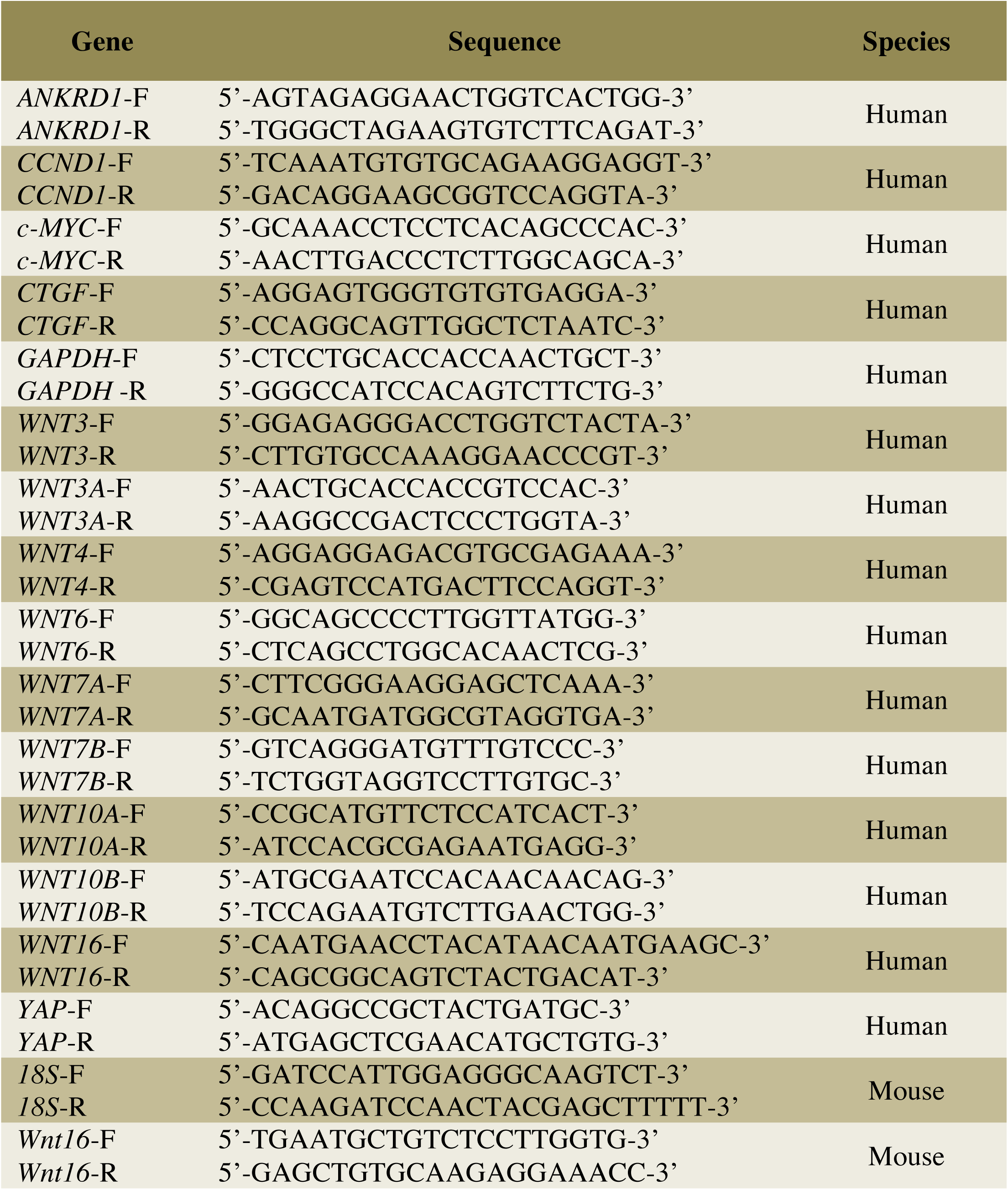
List of human and mouse primer sequences.

